# Metabolic responses at sub-thermoneutral temperatures in captive common waxbills (*Estrilda astrild*)

**DOI:** 10.1101/2025.10.28.685071

**Authors:** Cesare Pacioni, Hanne Danneels, Laura Dutrieux, Marina Sentís, Luc Lens, Diederik Strubbe

## Abstract

Understanding how endotherms manage energy expenditure at sub-thermoneutral temperatures is crucial for predicting their physiological flexibility and adaptive potential. This study investigated the metabolic responses of captive-bred common waxbills (*Estrilda astrild*) to decreasing ambient temperatures, with a focus on three questions: whether basal metabolic rate (BMR) predicts standard metabolic rate (SMR), how consistent individual metabolic responses are, and whether waxbills employ cold-induced energy-saving mechanisms. We found that BMR did not reliably predict SMR, underscoring the limits of BMR as a proxy under ecologically relevant conditions. SMR showed moderate repeatability across individuals, but variation was mainly due to baseline metabolic level rather than slope differences in thermal reaction norms. A two-breakpoint model best described the relationship between temperature and metabolic rate, with a marked decline in SMR below ~18.8°C. This decrease, along with observed reductions in core body temperature in some individuals, reveals an energy-saving strategy involving shallow hypothermia at relatively mild sub-thermoneutral temperatures. These findings suggest that energy-saving mechanisms may be intrinsic to the species rather than purely plastic responses. Overall, our results highlight both the limitations of BMR and the importance of measuring metabolic responses across thermal gradients to detect subtle but ecologically relevant hypometabolic strategies.

## 1. Introduction

Metabolic rate is an important physiological trait that underpins energy balance, survival and reproductive success in endothermic animals (McNab, 2012). As energy resources are finite, individuals must allocate energy strategically across life-history demands to optimize fitness (Burger et al., 2019; Stearns, 1992). The basal metabolic rate (BMR) represents a standardized measure of the minimum energy required for maintenance within the thermoneutral zone (TNZ; McNab, 2012), and it can constitute a substantial share of total daily energy expenditure (DEE). The TNZ is defined as the range of ambient temperatures (lower and upper critical temperatures) within which an animal can maintain its body temperature without altering metabolic heat production or water loss via evaporation (IUPS Thermal Commission, 2001). The relationship between metabolic rate and ambient temperature is typically explained by the Scholander-Irving model of endothermic homeothermy (Scholander et al., 1950). Outside the TNZ, metabolic rate increases linearly above BMR to maintain normothermia. Given its crucial role in maintaining energy balance, BMR is often considered a reliable indicator of an animal’s overall energy demands and it has been linked to various biological processes, e.g. thermoregulation, activity, reproduction, or other energy-demanding activities (Biro & Stamps, 2010; Burton et al., 2011; Glazier, 2005, 2015; Mckechnie & Swanson, 2010; McNab, 1997; Speakman et al., 2004). Moreover, it shows moderate repeatability, but its stability varies across contexts and declines over time, reflecting both individual consistency and environmental influence (Auer et al., 2016).

However, despite its widespread use, the extent to which BMR reflects overall DEE remains debated (Daan et al., 1990; Ricklefs et al., 1996; Visser et al., 2019). Intraspecific studies that measure both BMR and DEE in the same individuals are still relatively rare, and their findings have been inconsistent. Some studies report a positive correlation between BMR and DEE (Nilsson, 2002; Rezende et al., 2009; Tieleman et al., 2004), suggesting that BMR may serve as a useful proxy for overall energy turnover. Others have found no such relationship after accounting for intrinsic (e.g., age, body size, and reproductive status) and extrinsic factors (e.g., temperature and food availability; Fyhn et al., 2001; Meerlo et al., 1997; Peterson et al., 1998; Speakman et al., 2003). For instance, in captive zebra finches (*Taeniopygia castanotis*), BMR was positively associated with DEE outside the breeding season, but this correlation disappeared during reproduction, illustrating how ecological and life-history context can modulate the BMR–DEE relationship (Careau et al., 2013; Vézina et al., 2006). Briga and Verhulst (2017) further highlighted the limitations of BMR as a predictor by comparing measurements within and outside the thermoneutral zone. They found that, although BMR accounts for a substantial share of energy expenditure (~30%; see also Careau et al., 2013; Daan et al., 1990), it did not predict metabolic rates at lower ambient temperatures, where additional thermoregulatory costs arise. This suggests that while BMR is informative under standardized conditions, it may not capture the energetic demands animals face in more variable thermal environments.

While the Scholander–Irving model provides a useful baseline, individuals may differ in how sharply their metabolic rate increases outside the TNZ. Such variation is often described using reaction norms, which capture how phenotypic traits change across environmental gradients for different genotypes or individuals. In this framework, some individuals may exhibit a steep rise in metabolic rate as temperatures fall, while others show a shallower increase, reflecting consistent differences in energetic responses to cold (Briga & Verhulst, 2017). Beyond differences in slope, some species deviate from the expected linear increase altogether by reducing metabolic output at low temperatures. Such energy-saving strategies include torpor in hummingbirds, where animals lower body temperature and metabolic rate at night instead of sustaining the high costs predicted by extrapolation of the Scholander–Irving curve (Shankar et al., 2022). More subtle forms of hypometabolism, such as nocturnal hypothermia (Carr & Lima, 2013), have been hypothesized in other species, though their prevalence and ecological significance remain debated (Fristoe et al., 2015; Ritchison, 2023; Zhang et al., 2018). For example, an interpopulation comparison of great tits (*Parus major*) showed that at –10 °C, metabolic rates were lower than expected, indicating that both northern (Finland) and southern (Sweden) birds entered controlled hypothermia at night (Broggi et al., 2004). The authors suggest that their finding may indicate that nocturnal hypothermia may be a widespread but underappreciated strategy in small passerines to reduce energetic costs during winter nights. Such physiological adjustments may also play a role in shaping invasion success of introduced species, as species confronted with novel climates may persist or fail depending on their capacity to manage energetic costs under thermal stress (Davidson et al., 2011; Strubbe et al., 2015, 2023).

The common waxbill (*Estrilda astrild*) is a highly successful invasive species, known for its broad and expanding distribution beyond its native range. Originally from sub-Saharan Africa, this estrildid finch has colonized a wide range of environments, including the Iberian Peninsula (Reino, 2005) and numerous tropical and subtropical islands (Stiels et al., 2011). Its native range already spans diverse climatic regions (i.e., from the Mediterranean conditions of South Africa to the humid tropics of the Rift Valley and the arid Sahel), indicating a high degree of ecological flexibility. In Brazil, its spread has been attributed to a combination of intrinsic adaptability and extrinsic factors such as intentional releases, predator release, infrastructure corridors, and habitat modification (da Silva et al., 2018). Despite this ecological success, little is known about the species’ thermal physiology or how it manages energy demands across contrasting environments. Recent findings by Pacioni et al. (2024) showed that wild-caught waxbills in South Africa reduced their metabolic rate at lower ambient temperatures, deviating from predictions based on the Scholander–Irving model. This reduction may represent an energy-saving strategy, potentially enhancing survival in colder conditions and contributing to the species’ ability to establish in non-native temperate regions. Because such strategies typically involve coordinated decreases in both metabolic rate and core body temperature, examining body temperature responses alongside metabolic rate is essential to assess whether waxbills employ mechanisms akin to nocturnal hypothermia.

Similar downregulation of metabolic rates in response to cold has been observed in other bird species (Bush et al., 2008; Smit & McKechnie, 2010; Thabethe et al., 2013). For instance, Lovegrove (2000, 2003) suggested that while temperate-zone species often increase BMR in winter to meet thermogenic demands, birds from milder climates (e.g., Afrotropical region) tend to adopt energy conservation as their primary seasonal strategy. At the same time, evidence from temperate species such as great tits (*Parus major*) shows that controlled nocturnal hypothermia can also occur during cold conditions (Broggi et al., 2004). Taken together, these findings suggest that hypometabolic responses may occur across climatic zones, although the circumstances under which they are expressed remain uncertain and debated. Moreover, it is not clear to what extent such responses are in fact energetically beneficial: lowering metabolism usually entails lowering body temperature, which may increase predation risk because animals may be slower to detect and react to threats (Barratt et al., 2025), and birds that cool substantially at night also face the costs of reheating in the morning. How the balance of these costs and benefits plays out under natural conditions remains unresolved.

In this study, we examine the metabolic responses of captive-bred common waxbills (*Estrilda astrild*) at sub-thermoneutral temperatures to address three research questions: (i) does basal metabolic rate (BMR) predict metabolic rates at lower ambient temperatures?; (ii) how consistent are individual metabolic reaction norms to decreasing temperatures?; (iii) do individuals show metabolic rates at cold temperatures that deviate from the linear increases predicted by the Scholander–Irving model?; and (iv) is core body temperature maintained or reduced under low ambient temperatures? In contrast to previous work on wild waxbill populations that documented lower-than-expected metabolic rates at the population level (Sentís et al., 2025), our experimental design allows us to test these deviations at the level of individual birds. By focusing on individuals, we may be able to uncover variation in metabolic strategies that may be masked in population-level analyses, allowing for deeper insight into the physiological mechanisms underlying species’ responses to cold.

## 2. Material and methods

This study involved 12 captive-bred common waxbills, acquired from certified breeders in Belgium. From the time of arrival (October 2023) until the conclusion of the experiments (May 2024), the birds were housed in an indoor aviary measuring 200 × 100 × 200 cm (length × width × height). Food and water were provided ad libitum. The ambient temperature fluctuated around 19°C, with relative humidity ranging between 30% and 50% during the measurement period. All individuals were sexed (sexed based on their under-tail coverts, black for males and brown for females) and marked with unique combinations of three coloured plastic rings for visual identification. The experimental procedures were approved by the Ethical Committee for Animal Experimentation at Ghent University (IRC-UGent EC2023-110).

### 2.1 Metabolic measurements

Rest metabolic rate (RMR) was measured using open-flow respirometry (Lighton, 2018) to assess oxygen consumption (VO□) during the birds’ inactive phase. The protocol followed Pacioni et al. (2023), with slight adaptations. Measurements were conducted overnight, beginning between 17:00 and 19:00 and ending between 07:00 and 08:00 the following morning. Birds were placed individually into airtight plastic metabolic chambers (1.1 L), each equipped with a plastic mesh floor for support and comfort. Each chamber contained an air inlet at the top and an outlet near the bottom to ensure efficient mixing of incoming and outgoing air. Up to seven birds were measured per night, with an additional chamber left empty to record baseline values. Chambers were placed inside a darkened, temperature-controlled cabinet (PELT 5, Sable Systems), where ambient temperature was set. The temperatures tested ranged from 11 to 35.5 °C. One temperature was tested each night. Ambient air was pumped into each chamber at a flow rate of 650 mL/min, regulated by a mass flow controller (FB-8, Sable Systems), to maintain CO□ concentrations below 0.5%. Excurrent air was routed through a multiplexer (RM-8, Sable Systems), allowing sequential sampling of each chamber. The outcoming air was pulled through a Field Metabolic System (FMS-3, Sable Systems). Measurements were carried out in cycles throughout the night, resulting in approximately 14.5 hours of data collection per session. The first 2 hours of measurement were discarded to ensure birds were in a post-absorptive status (Pacioni et al., 2024). Each bird was weighed to the nearest 0.01 g immediately before and after each metabolic measurements. Core body temperature was recorded on a subset of individuals immediately following each respirometry measurements using a T-type thermocouple (Omega®) with a precision of 0.01 °C. The probe tip was coated and lubricated with Vaseline® to facilitate smooth insertion, and was gently inserted no more than one centimeter into the bird’s cloaca. Handling was kept under one minute to reduce stress, and temperature readings were continuously logged using the PicoLog TC-08 (Pico Technology, UK) interface and software.

### 2.2 Respirometry and data analysis

ExpeData software (Sable Systems) was used to record each experimental trial and extract rest metabolic rate values (mL O_2_/min). O_2_, CO_2_, and water vapor pressures were measured in both the incurrent and excurrent streams. To estimate rest metabolic rates, the lowest stable section of the curve (averaged over 5 min) was selected using equation 9.7 from Lighton (2018):

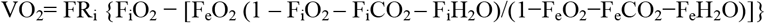

where VO_2_ represents the rate of oxygen consumption (ml O_2_/min), FR_i_ is the incurrent flow rate, F_i_ the fractional concentrations of the incurrent gas and F_e_ the fractional concentrations of the excurrent gas.

To model individual metabolic reaction norms across temperature, each metabolic rate measurement was assigned to a temperature category based on the ambient temperature at which it was recorded. For measurements obtained below the thermoneutral zone, we refer to metabolic rates as standard metabolic rate (SMR), following established usage in avian physiology (e.g., Briga & Verhulst, 2017). Following Briga & Verhulst (2017) and based on the outcome of the inflection point analysis (see below), empirically defined temperature ranges were used to classify observations into three categories: 28–35.5°C for BMR, 20–26°C for SMR_24_, and 12–18°C for SMR_16_. Individual metabolic reaction norms to sub-thermoneutral ambient temperatures were characterized using Bayesian linear mixed-effects models. The response variable was SMR, with ambient temperature and body mass included as fixed effects. Individual identity (ID) was included as a random grouping factor. To assess variation among individuals in their metabolic response to temperature (i.e., reaction norms), models included a random slope of ambient temperature nested within individuals (Briga & Verhulst, 2017). Model performance was compared between random slope and simpler random intercept models using leave-one-out cross-validation (LOO) and Widely Applicable Information Criterion (WAIC). Bayes factors were also computed to quantify the relative support for each model.

The relationship between rest metabolic rates and ambient temperature was evaluated using a generalized additive mixed model (GAMM), incorporating individual identity as a random effect to account for repeated measures (implemented via the gamm function in the ‘mgcv’ package; Wood, 2023). To detect potential inflection points in this relationship, a segmented regression analysis was performed using the ‘segmented’ package in R (Muggeo, 2008). The overall shape of the curve was visualized with a GAMM smoother. Models with one and two breakpoints were compared based on the corrected Akaike Information Criterion (AICc). Repeatability of BMR and SMR was estimated using the rpt function from the rptR package (Stoffel et al., 2017). Linear mixed-effects models were fitted with log-transformed BMR or SMR as the response variable and ambient temperature as a fixed effect. Individual identity (ID) was included as a random intercept to partition within- and between-individual variance components. Repeatability was calculated as the proportion of total variance explained by ID, and uncertainty was assessed using 1000 parametric bootstraps and 1000 permutations.

To assess the relationship between ambient temperature and core body temperature, the analysis focused on a subset of individuals measured across a gradient of three ambient temperatures (11°C, 22°C, and 30°C). These temperatures were selected to cover a range from below to above the birds’ thermoneutral zone, allowing to test whether core body temperature is actively maintained or allowed to decrease at low ambient temperatures. Ambient temperatures were chosen independently of the SMR trials (16□°C and 24□°C) to specifically capture the response of core temperature across a broader thermal gradient. A linear mixed-effects model was used, with core body temperature as the dependent variable, ambient temperature (categorical) as a fixed effect, and ID included as a random intercept to account for repeated measures within individuals.

Analyses were conducted using both whole-body metabolic rates and values adjusted for body mass. Sex was included as an independent factor in all the models. Mass-independent metabolic rates were calculated as residuals from linear regressions of log-transformed rest metabolic rates against log-transformed body mass. An alternative method, incorporating log body mass as a covariate within the models, was also tested. As both approaches yielded consistent outcomes, the residual method was adopted for presenting mass-independent metabolic rates in this study. The level of significance was set at p < 0.05. The body mass and rest metabolic rates were log10-transformed before the analysis. All statistical analyses were performed using R software v. 4.2.2 (R Core Team 2022).

## 3. Results

To address question (i), we tested whether BMR predicts metabolic rates at lower ambient temperatures. Correlations between whole-body BMR (n = 68) and both SMR16 (n = 38) and SMR24 (n=45) were weak and not statistically significant (r = 0.32, p = 0.334; r = 0.40, p = 0.327 respectively). By contrast, SMR16 and SMR24 were strongly correlated with each other (r = 0.96, p < 0.001), indicating consistency in SMR across cold treatments. Similar results were found for mass-independent values. To address question (ii), we examined the consistency of individual metabolic reaction norms (n = 11) across sub-thermoneutral temperatures. The random slope model showed only marginal improvement in fit compared to the intercept-only model (WAIC = −105.3 vs. −103.6; LOO = −104.0 vs. −101.5), while Bayes factor analysis strongly supported the simpler model (BF ≈ 1015). Thus, models indicated variation in baseline metabolic rates among individuals (random intercept SD ≈ 0.06), but not in their slopes (SD ≈ 0.00; Figure 1). Repeatability estimates revealed moderate individual consistency for metabolic traits. Whole-body BMR repeatability was 0.33 (SE = 0.14; 95% CI = 0.03–0.58; p < 0.001), whereas mass-independent BMR had lower, marginally non-significant repeatability (0.21; SE = 0.14; 95% CI = 0.00–0.49; p = 0.066). Whole-body SMR showed higher repeatability (0.43; SE = 0.15; 95% CI = 0.08–0.66; p < 0.001) and mass-independent SMR was similarly repeatable (0.31; SE = 0.14; 95% CI = 0.00–0.57; p < 0.001). To address question (iii), we used GAMMs and inflection point analyses to test whether individual metabolic rates (n=151) at cold ambient temperatures deviated from the linear increases predicted by the Scholander–Irving model. The analyses identified two change points: the first at 28.9□± 1.2 °C, corresponds to the lower critical temperature (LCT), below which metabolism begins to increase as temperature decreases. The second at 18.8□± 1.7 °C, reflects a deviation from the expected linear response (i.e., Scholander– Irving model; Figure 2). To address question (iv), we analyzed how core body temperature varied across different ambient temperatures and found that core body temperature was positively correlated with ambient temperature (Spearman’s ρ = 0.50, p = 0.036). On average, individuals had higher core body temperatures at higher ambient temperatures. Birds at 30°C had the highest mean core body temperature (41.48 ± 0.56 °C, n = 6), while birds at 12°C and 22°C had slightly lower temperatures (40.37 ± 0.92 °C, n = 7, and 40.01 ± 1.38 °C, n = 4, respectively).

**Figure 1.**
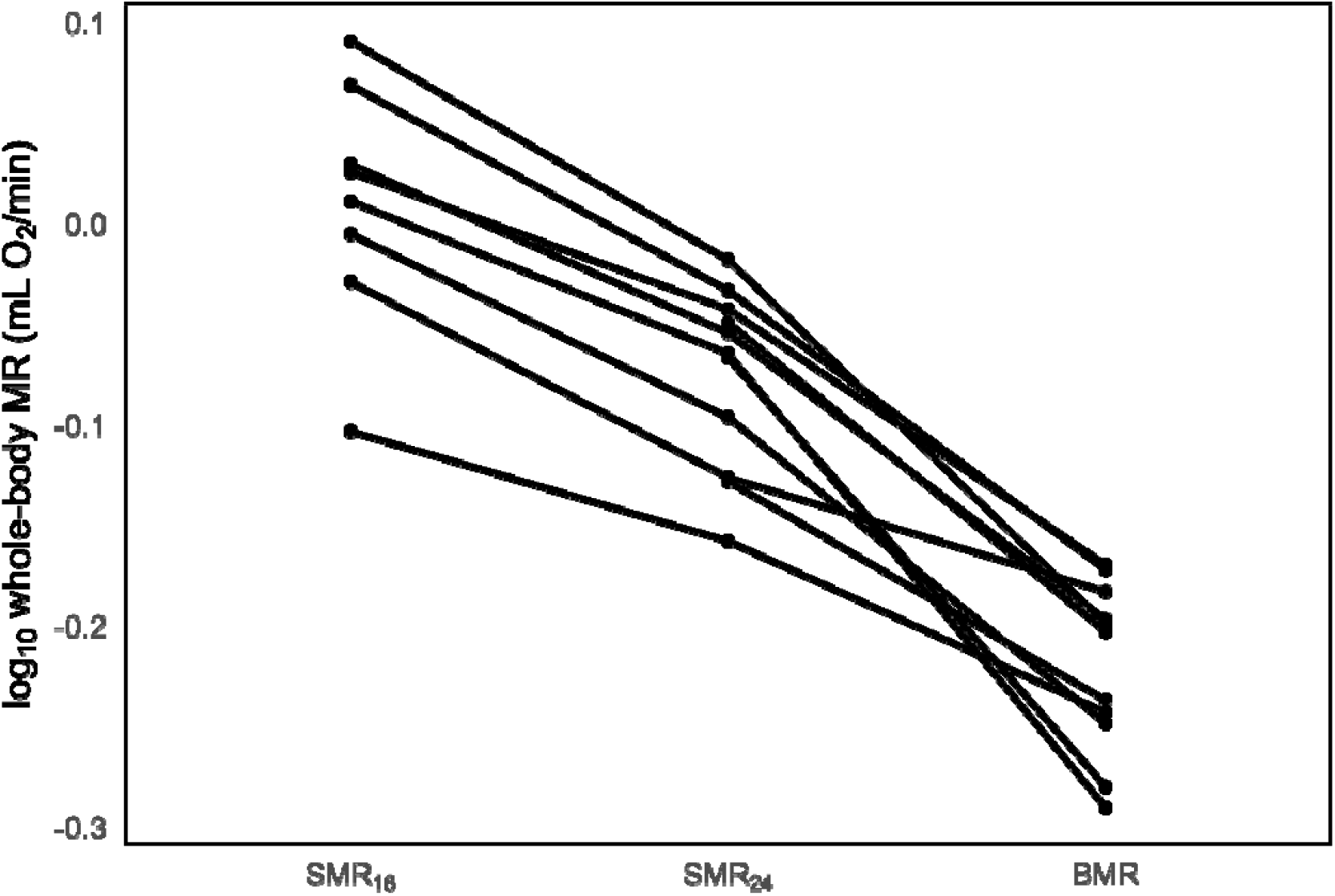
Individual (mean) reaction norms for common waxbills (*Estrilda astrild*) with complete measurements across all three metabolic rate categories (SMR_16_, SMR_24_, and BMR).

**Figure 2.**
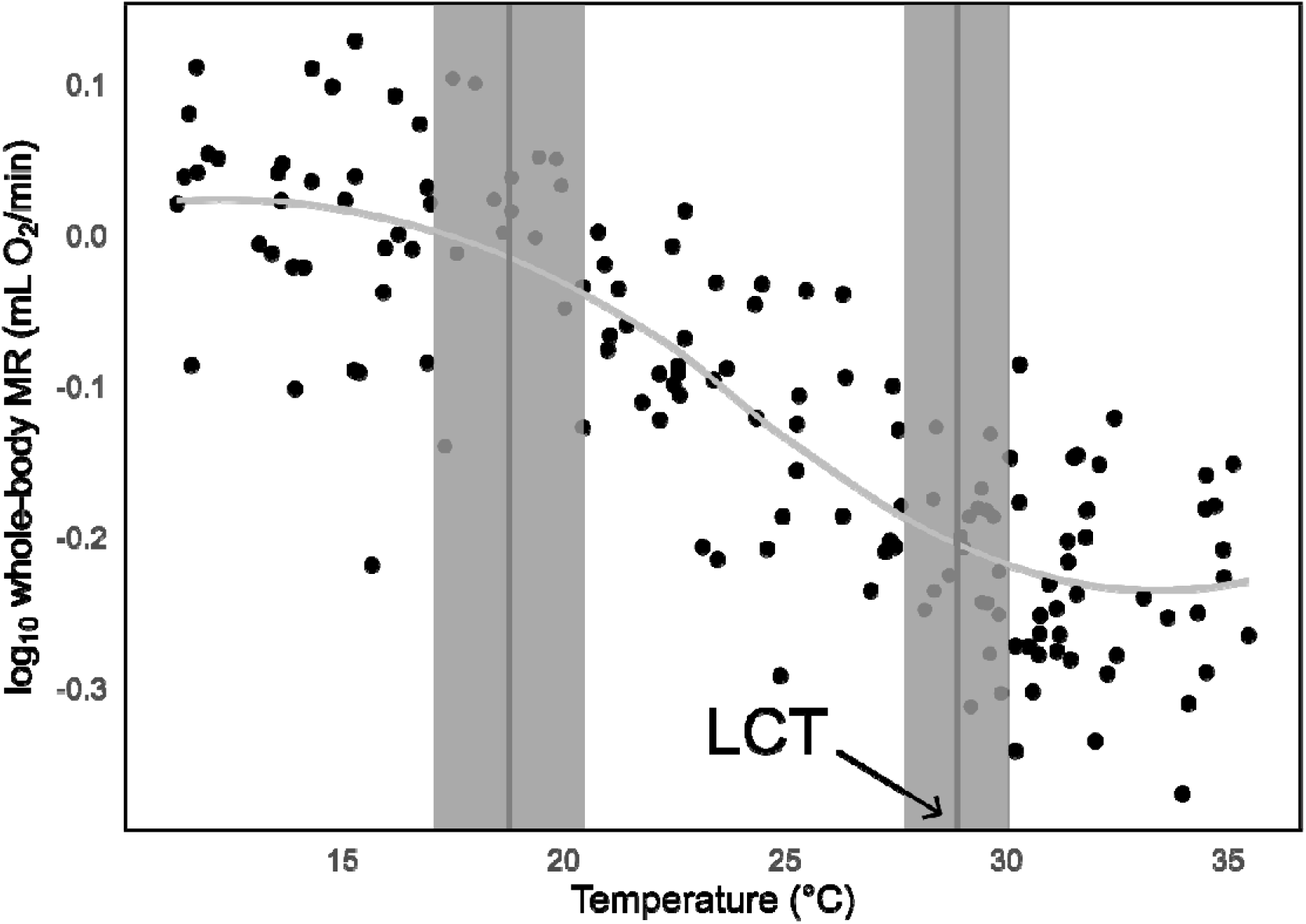
Log10 whole-body metabolic rate (MR; mL O_2_/min; n = 151) of captive common waxbills (*Estrilda astrild*) measured across a range of ambient temperatures (°C). The two vertical lines indicate inflection points at 18.8 ± 1.7 °C and 28.9 ± 1.2 °C (lower critical temperature; LCT). Light grey bands show ± standard deviation around each inflection point.

## 4. Discussion

This study examined the metabolic responses of captive-bred common waxbills to sub-thermoneutral temperatures, focusing on whether resting metabolism can predict energy use at colder temperatures, the consistency of individual metabolic responses, potential deviations from the Scholander–Irving model, and changes in core body temperature. To highlight our most novel result, we begin with research question iii). The two-breakpoint model best described the relationship between ambient air temperature and metabolic rate, indicating a lower critical temperature at ~28.9 °C and an additional inflection at ~18.8 °C. Below this threshold, metabolic rates no longer increased linearly as predicted by the Scholander–Irving model but levelled off. This deviation, together with modest reductions in body temperature, suggests a shallow hypothermia strategy that allows waxbills to conserve energy at relatively mild sub-thermoneutral temperatures. Rather than reflecting a metabolic ceiling, this pattern points to an intrinsic energy-saving mechanism that may contribute to ecological flexibility and invasive success.

Our third research question asked whether common waxbills deviate from the linear increases in metabolic rate predicted by the Scholander–Irving model. We identified a lower critical temperature of 28.9 °C, consistent with values reported previously in a smaller sample of captive-bred common waxbills (Pacioni et al., 2023), but here confirmed with a larger dataset. Below this threshold, metabolic rates initially rose as expected with decreasing ambient temperature, reflecting the increased energetic demands of thermoregulation. However, at ~18.8 °C we detected a distinct inflection point, after which metabolic rates no longer followed the linear increase predicted by the Scholander–Irving model but instead leveled off. A comparable pattern was also observed in wild South African waxbills (Pacioni et al., 2024), though in that study different individuals were measured at each temperature and wider increments (4 °C) were used, making it more difficult to resolve fine-scale patterns. By contrast, our repeated-measures design under stable laboratory conditions provides stronger evidence that this non-linear response possibly represents a consistent feature of waxbill physiology. Importantly, this deviation is unlikely to reflect a metabolic ceiling: summit metabolic rates in common waxbills have been estimated at 1.75–3.47 mL O□/min (Pacioni et al., 2023), whereas in our study overnight rates never exceeded 2.0 mL O□/min. Although part of this discrepancy may be explained by diurnal effects (night-time metabolism is typically 20–25% lower than during the day), this alone cannot account for the reduced values. Thus, the birds in our study were clearly not operating at or near their maximal metabolic capacity, even under the coldest conditions tested. Notably, the deviation occurs at relatively mild sub-thermoneutral temperatures (~16–18 °C), whereas other passerines such as e.g. Scandinavian great tits allow body temperature to decline only at much lower ambient temperatures (Broggi et al., 2004).

The observed deviation from Scholander–Irving predictions therefore possibly reflects an energy-saving strategy rather than a metabolic limitation. Instead of continually elevating metabolic output as temperatures decline, waxbills appear to conserve energy by limiting further increases at relatively mild sub-thermoneutral conditions. Our measurements of body temperature, although limited, provide additional support: while birds maintained the highest core temperatures at 30 °C (mean 41.48 °C), very close to the ~42 °C generally expected for small passerines under thermoneutral conditions, they showed a modest but consistent reduction already at 22 °C (mean 40.01 °C) and a similar reduction at 11 °C (mean 40.37 °C). Such shallow hypothermia may represent a controlled adjustment that helps lower thermoregulatory costs without incurring the risks of torpor. A recurrent criticism of this interpretation is that lowering body temperature may also entail costs, including slower responsiveness and elevated predation risk (Carr & Lima, 2013), as well as the energetic burden of reheating once temperatures rise. Moreover, theoretical work suggests that the relative benefits of hypothermia diminish under severe cold, where rewarming costs can outweigh savings (Brodin et al., 2017). However, our finding that waxbills reduce body temperature already at relatively mild sub-thermoneutral temperatures (~22 °C) may mitigate this concern, as the smaller gradient between internal and external temperatures means that modest body temperature reductions are more likely to yield net energy savings. Similar shallow hypothermia has been documented in Northern Cardinals (*Cardinalis cardinalis*), which reduced body temperature by ~1.3 °C, leading to estimated overnight energy savings of 7–16% (Schaeffer et al., 2015), a magnitude comparable to the ~1 °C reduction we observed in waxbills. Taken together, these lines of evidence suggest that common waxbills combine modest adjustments in body temperature with reduced increases in metabolic output to limit energy expenditure when exposed to temperatures below their TNZ.

Regarding our first research question, Briga and Verhulst (2017) examined the relationship between BMR and SMR in zebra finches and showed that while SMRs at different sub-thermoneutral temperatures were strongly correlated, BMR was only weakly associated with SMR. They traced this weak link to differences in body temperature regulation: some individuals maintained a high body temperature at low ambient temperatures and showed steep increases in metabolic rate, whereas others allowed body temperature to drop and showed shallower slopes. This heterogeneity in thermoregulatory strategy weakened the expected correlation between BMR and SMR. The weak coupling is also consistent with their argument that BMR and SMR are underpinned by partly different physiological determinants: BMR reflects primarily central organ size and function, whereas SMR additionally depends on insulation, thermogenic capacity, and body temperature regulation. In our waxbill study, we found a similar pattern: BMR did not correlate with SMR, while SMRs were strongly correlated with each other. Our data therefore support the conclusion that BMR is a poor predictor of metabolic performance at sub-thermoneutral temperatures. Unlike Briga and Verhulst (2017), we could not test directly whether variation in body temperature regulation explains this decoupling, because we only measured body temperature in a subset of individuals and not systematically across all temperature treatments. Nonetheless, the parallels with the zebra finch results suggest that body temperature regulation may likewise play a role in shaping the weak association between BMR and SMR in common waxbills. These findings underscore the importance of measuring metabolic rate not only within, but also outside the thermoneutral zone, especially at temperatures that birds are likely to encounter frequently during their daily and seasonal cycles.

Our second research question examined how consistent individual metabolic reaction norms are to decreasing ambient temperatures. Briga and Verhulst (2017) showed that zebra finches varied in the slopes of their reaction norms, with pronounced differences under favourable conditions but reduced variation when conditions were harsher, suggesting that environmental context can constrain individual flexibility. Similar patterns have been reported in ectotherms: juvenile barramundi fish (*Lates calcarifer*) showed pronounced slope variation in metabolic traits under benign conditions, with predictable but divergent responses to environmental stressors (Norin et al., 2016). Our waxbills align more closely with the latter result, showing no evidence of slope variation: individuals differed in their baseline metabolic level but responded to declining ambient air temperatures with a shared rate of increase. Because the birds were housed under stable aviary conditions with ad libitum food, it seems unlikely that environmental constraints explain this uniformity. More plausible explanations are that waxbills follow a more uniform thermoregulatory strategy than zebra finches, or that limitations in sample size reduced our ability to detect slope differences.

Future research on avian thermoregulatory patterns would benefit from integrating metabolic and body temperature measurements with spatially explicit physiological modelling. While our study highlights the need for continuous monitoring of core body temperature alongside metabolic rates, a broader challenge is to assess whether shallow hypothermia confers ecological advantages under natural conditions. Biophysical approaches (Briscoe et al., 2023; Sentís et al., 2023) can link species’ morphological and physiological traits with climatic and microhabitat variation, and generate predictions of energy expenditure under different scenarios, for instance, those based on classical Scholander–Irving expectations versus the deviations observed here. Such models can be used to produce spatially explicit maps of predicted energetic costs, which in turn can guide more targeted empirical sampling by identifying populations and environments that are most informative for testing hypotheses about energy use. Applying this model-guided approach to common waxbills would allow more definitive evaluation of whether the patterns we observed translate into ecological benefits, and whether they may contribute to the species’ capacity to invade and expand in non-native environments.

## 5. Conclusions

Our results demonstrate three main points. First, BMR is not a reliable predictor of SMR under sub-thermoneutral conditions. Second, metabolic rates showed moderate repeatability, but individual differences were mainly in baseline levels rather than in slopes of thermal reaction norms. Third, waxbills did not continue to increase metabolism linearly below ~18.8 °C. Instead, they displayed a distinct levelling off of metabolic rate, coupled with modest body temperature reductions. This pattern strongly suggests an intrinsic energy-saving strategy involving shallow hypothermia. By limiting further increases in metabolic output at relatively mild sub-thermoneutral temperatures, common waxbills may reduce energetic costs in ways that facilitate persistence in novel environments and may contribute to their invasive success. Future studies combining continuous body-temperature monitoring with spatially explicit physiological modelling will be crucial to test whether these patterns translate into ecological advantages under natural conditions.

## Data and code availability

All datasets and R script used in this study are available as supplementary material and have been archived on Figshare at https://figshare.com/s/36c03c287f076ea8cd97

## CRediT authorship contribution statement

**Cesare Pacioni:** Conceptualization, Methodology, Formal analysis, Investigation, Visualization, Writing – original draft. **Hanne Danneels:** Data curation, Investigation, Formal analysis, Writing – review & editing. **Laura Dutrieux:** Data curation, Investigation, Formal analysis, Writing – review & editing. **Marina Sentís:** Investigation, Writing – review & editing. **Luc Lens:** Conceptualization, Supervision, Funding acquisition, Writing – review & editing. **Diederik Strubbe:** Conceptualization, Methodology, Supervision, Writing – review & editing.

